# Neural correlates of spatiotemporal properties of lightness induction

**DOI:** 10.1101/2023.07.28.550941

**Authors:** Amna Malik, Huseyin Boyaci

## Abstract

The lightness of a surface depends not only on the amount of light reflected off it but also on the context it is embedded in. Despite a long history of research, neural correlates of context-dependent lightness perception remain a question of ongoing debate. The current study seeks to expand upon the existing literature by measuring fMRI responses to a dynamic version of a classical effect called simultaneous lightness induction (SLI), where a central disk appears lighter when embedded in a darker surround compared to when embedded in a lighter surround. During the fMRI experiment, we presented ten participants with a dynamic SLI stimulus by modulating the luminance of either the achromatic surround (surround-modulation condition) or the achromatic disk (disk-modulation condition) at four different frequencies ranging from 1 to 8 Hz. Behaviorally, when the surround luminance is modulated at low frequencies, participants perceive an illusory change in the lightness of the disk (lightness induction). In contrast, they perceive little or no induction at higher frequencies. Utilizing this temporal dependence and controlling for long-range responses to border contrast and luminance changes, we found that activity in the primary visual cortex (V1) correlates with lightness induction. However, such a correlation was not evident in extrastriate areas, V2, V3, and V4. These findings provide evidence for the involvement of V1 in the processing of context-dependent lightness.

## Introduction

Lightness is defined as the perceived relative reflectance of a surface, and it depends not only on the amount of light reflected off the surface but also on its context. For example, in the well-known simultaneous lightness induction (SLI) effect (Figure 1, also known as simultaneous brightness contrast), a mid-grey patch is perceived as lighter when embedded in a darker surround than when embedded in a lighter surround (Alhazen, 1883; Heinemann, 1955; Kingdom, 1997; Von Helmholtz, 1867). Such context-dependent effects are frequently used in the literature to study lightness perception and its neural correlates because they allow researchers to dissociate lightness and luminance (Adelson, 1993; Blakeslee & McCourt, 1997; Cornelissen, Wade, Vladusich, Dougherty, & Wandell, 2006; Haynes, Lotto, & Rees, 2004; Komatsu, 2006; Pereverzeva & Murray, 2008; Rossi & Paradiso, 1996, 1999; Sinha et al., 2020). Results of those and other studies in the literature, however, have not yet led to a consensus about the visual areas and neural mechanisms involved in the processing of lightness. The proposed mechanisms can be broadly grouped into three categories; a low-level account (Anderson, Dakin, & Rees, 2009; Blakeslee & McCourt, 1999; Dakin & Bex, 2003), a mid-level account (Bouvier, Cardinal, & Engel, 2008; Boyaci, Fang, Murray, & Kersten, 2010; Economou, Zdravkovic, & Gilchrist, 2007; Gilchrist et al., 1999) and a high-level account (Adelson, 1993; Perna, Tosetti, Montanaro, & Morrone, 2005). Low-level accounts argue that early retinotopic visual areas play an instrumental role in the process. In several studies, neural activity in those areas was indeed found to correlate with context-dependent lightness, providing evidence in favor of low-level accounts (Boyaci, Fang, Murray, & Kersten, 2007; Pereverzeva & Murray, 2008; Rossi, Rittenhouse, & Paradiso, 1996; Sasaki & Watanabe, 2004). Some other studies, however, provided opposite results (Cornelissen et al., 2006; Perna et al., 2005). Thus, whether and how low-level visual areas, including the primary visual cortex (V1), contribute to lightness processing remains an important open question.

**Figure 1:**
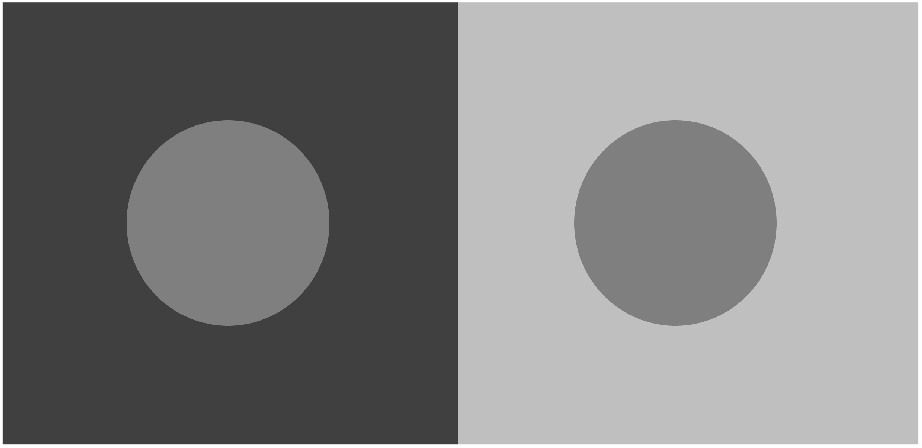
Simultaneous lightness induction (SLI). The disks have the same luminance, but the one on the darker background appears lighter than the one on the lighter background.

To answer this question, a dynamic version of the SLI effect, where a constant-luminance disk is perceived to vary in lightness when presented in a surround that is temporally modulated in luminance, is frequently used. A number of those studies have shown that neural activity in early retinotopic areas correlates with perceived lightness (e.g., Rossi & Paradiso, 1999.) Notably, however, in a human fMRI study using the dynamic SLI, Cornelissen et al. (2006) have found no evidence of context-dependent lightness processing in V1. In that study, authors recorded V1 responses to a foveally presented static disk embedded in a surround whose luminance varied at a relatively low frequency (1 Hz). They found that, in line with the SLI effect, the perceived lightness of the disk varied as a function of time. When they analyzed the fMRI BOLD responses in regions of V1 that retinotopically process the part of the visual field occupied by the disk, however, they found that the activity could be explained simply by long-range responses to the contrast variation at the border between the disk and the surround. Thus, they argued that there is no evidence of fMRI response to context-dependent lightness variations in V1 (Cornelissen et al., 2006).

Later, Pereverzeva and Murray (2008) re-examined the V1 activity using the dynamic SLI and a clever design that allowed dissociating responses to lightness and border contrast variations. Specifically, they presented the center disk at different luminances, ranging from very low to mid values. Perceptually the SLI effect increased as the disk luminance increased, whereas temporally integrated border contrast energy decreased. Results showed that in regions of V1 that retinotopically correspond to the disk’s center, there was a larger fMRI activity on trials where the central disk had higher luminance; in other words, the activity correlated with SLI, not border contrast. Thus, the study provided evidence that human V1 is involved in processing context-dependent lightness (Pereverzeva & Murray, 2008). In their design, however, the luminance of the central disk varied between different conditions. This could have confounded the results because the increase in the perceptual lightness effect was accompanied by an increase in the mean luminance, and an increase in mean luminance, independent of contrast, may increase the neural responses in early visual areas (Geisler, Albrecht, & Crane, 2007; Vinke & Ling, 2020).

Here, we followed a similar strategy to dissociate responses to context-dependent lightness and border contrast without changing the luminance of the central disk. To do so, we kept the central disk luminance constant and used different surround modulation frequencies ranging from low to high values, and collected fMRI BOLD responses (‘surround-modulation’ condition). The dynamic SLI effect decreases with frequency (De Valois, Webster, De Valois, & Lingelbach, 1986; Pereverzeva & Bromfield, 2013; Rossi & Paradiso, 1996), while border energy computed by integrating the contrast variation over time increases. Thus, we reasoned that this design should allow us to dissociate the responses to context-dependent lightness and border contrast and address the confound put forward by Cornelissen et al. (2006) and others. Further, in order to compare responses to luminance-dependent and context-dependent lightness variations, we have introduced a ‘disk-modulation’ condition in which the surround remained static, and the disk varied in luminance with the same temporal modulation characteristics as in the surround-modulation condition. Because the temporal contrast energy at the border for a particular frequency was equal across the two conditions, the difference between disk-modulation and surround-modulation conditions should reflect mainly the difference between neural responses to luminance and context-dependent lightness variations.

We noted that surround modulation alone might drive the fMRI responses without any lightness-related neural activity. This could happen because the fMRI response has a wide point-spread function and reflects the activity of a large number of functionally heterogeneous neurons in its spatial unit, i.e., a voxel. Even in a functionally identified cortical region of interest that retinotopically corresponds to the inside of the disk, there might be neural responses to the surround modulation or the overall illumination variation in the environment. To address this confound, we used a separate ‘control’ condition in which surround luminance varied as in the surround-modulation condition, but the disk was black, causing no lightness induction. Thus, comparing the surround-modulation and control conditions should allow us to address the possible confound of long-range luminance responses.

Our protocol, shown in Figure 2, included first a behavioral experiment to ensure that induction decreases with surround modulation frequency. As expected, the induction effect indeed decreased with frequency, consistent with findings in the literature. Next, in a block-design fMRI experiment, we collected responses under the surround-modulation, disk-modulation, and control conditions. We compared responses to these conditions in retinotopically identified regions of interest in the primary visual cortex (V1), as well as V2, V3, and V4 (Figure 4). If a region is involved in context-dependent lightness processing, we expect to find a strong response under the surround-modulation condition, approximately equal to that under the disk-modulation condition at low frequencies but not at high frequencies. Further, we expect surround-modulation condition responses to be higher than the control condition at low frequencies. If, however, any of these conditions are not satisfied, for example, if the responses under the surround-modulation condition do not decrease with frequency or if the responses at low frequencies do not differ between the surround-modulation and control conditions, that would call into question the involvement of that area in context-dependent lightness processing.

**Figure 2:**
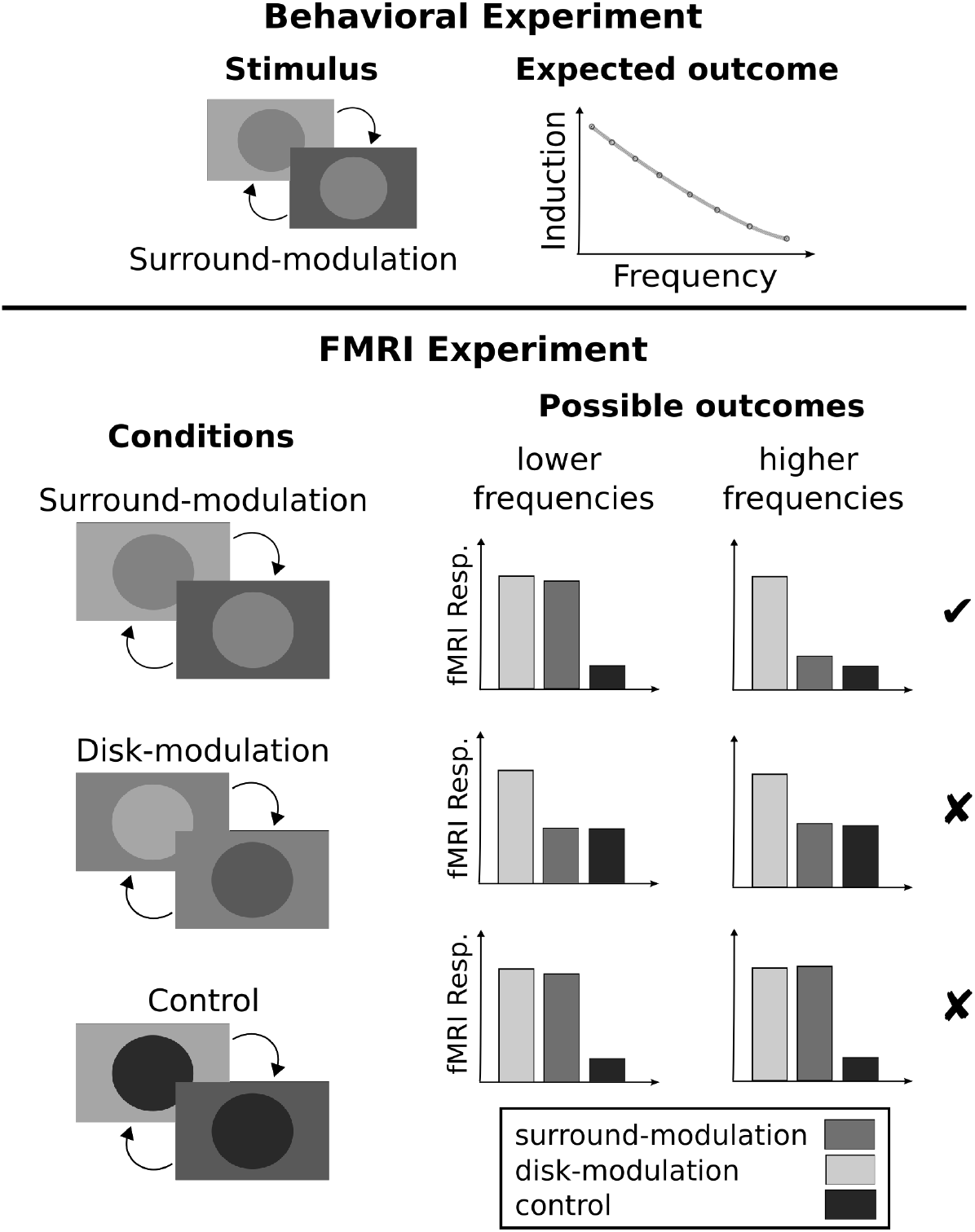
First, in a behavioral experiment, we measured the magnitude of induction using the dynamic version of SLI. Based on the results in the literature, we expect that induction decreases with the temporal frequency of the surround modulation. Next, in an fMRI experiment, we measured the fMRI responses under three stimulus conditions and at different modulation frequencies. The ‘surround modulation’ condition is similar to the stimulus in the behavioral experiment, which leads to the dynamic SLI effect at low but not high frequencies. In the ‘disk-modulation’ condition, the surround remains constant, and the disk luminance is modulated. In the ‘control’ condition, the surround is modulated as in the surround-modulation condition, but the disk is black, leading to no dynamic SLI effect. The right panel shows the possible outcomes. If a cortical region is involved in context-dependent lightness processing, the fMRI responses under the surround-modulation condition at low frequencies would be larger than those at high frequencies. Further, the responses under the surround-modulation condition would be larger than those under the control condition at low frequencies. The first row shows this possible outcome. Any outcome inconsistent with this pattern would call into question the involvement of that area in context-dependent lightness processing. The next two rows show two such possible outcomes. The first one would be from a region whose activity is not correlated with lightness and, at best, directly responds to luminance modulations in the surround. The bottom row shows the outcome if the region responds to the border contrast variation, not to the lightness variations in the disk.

## Materials and Methods

### Participants

Seven participants participated in the behavioral experiment, and a different set of ten participants participated in the fMRI experiment. All participants had normal or corrected-to-normal vision, and they gave their written informed consent before the experiments. Experimental procedures were approved by the Research Ethics Committee of Bilkent University.

### Behavioral Experiment

This experiment was conducted to measure the magnitude of dynamic SLI with different surround modulation frequencies and at different eccentricities within the central disk.

#### Experimental Setup and Stimulus

Visual stimuli were presented on a color reference LCD screen in an otherwise dark room (Eizo CG2730, size: 27 inches, pixel resolution: 2560 x 1440, refresh rate: 60 Hz). A luminance look-up table was prepared through direct measurement to characterize the monitor. The maximum and minimum achromatic luminance of the screen was 100 and 0.015 cd/m^2^, respectively. Participants were seated 60 cm from the screen with their head movements constrained using a head-and-chin rest. Participant responses were collected through a standard computer keyboard. Psychtoolbox functions (Brainard, 1997) on MATLAB (version 2018a, MathWorks Inc., New York, NY, USA) were used to program the experiments on a Dell 3630 Workstation running Ubuntu Linux (version 18.04).

Figure 3 shows the details of the experimental protocol. The basic stimulus consisted of an achromatic disk with a diameter of 11 degrees of visual angle, embedded in an achromatic rectangular surround subtending 23.54 by 13.37 degrees of visual angle. The rest of the screen was kept at minimum luminance. The luminance of the disk and surround was defined in terms of instrument luminance (IL)

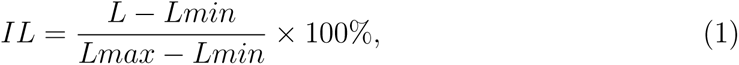

**Figure 3:**
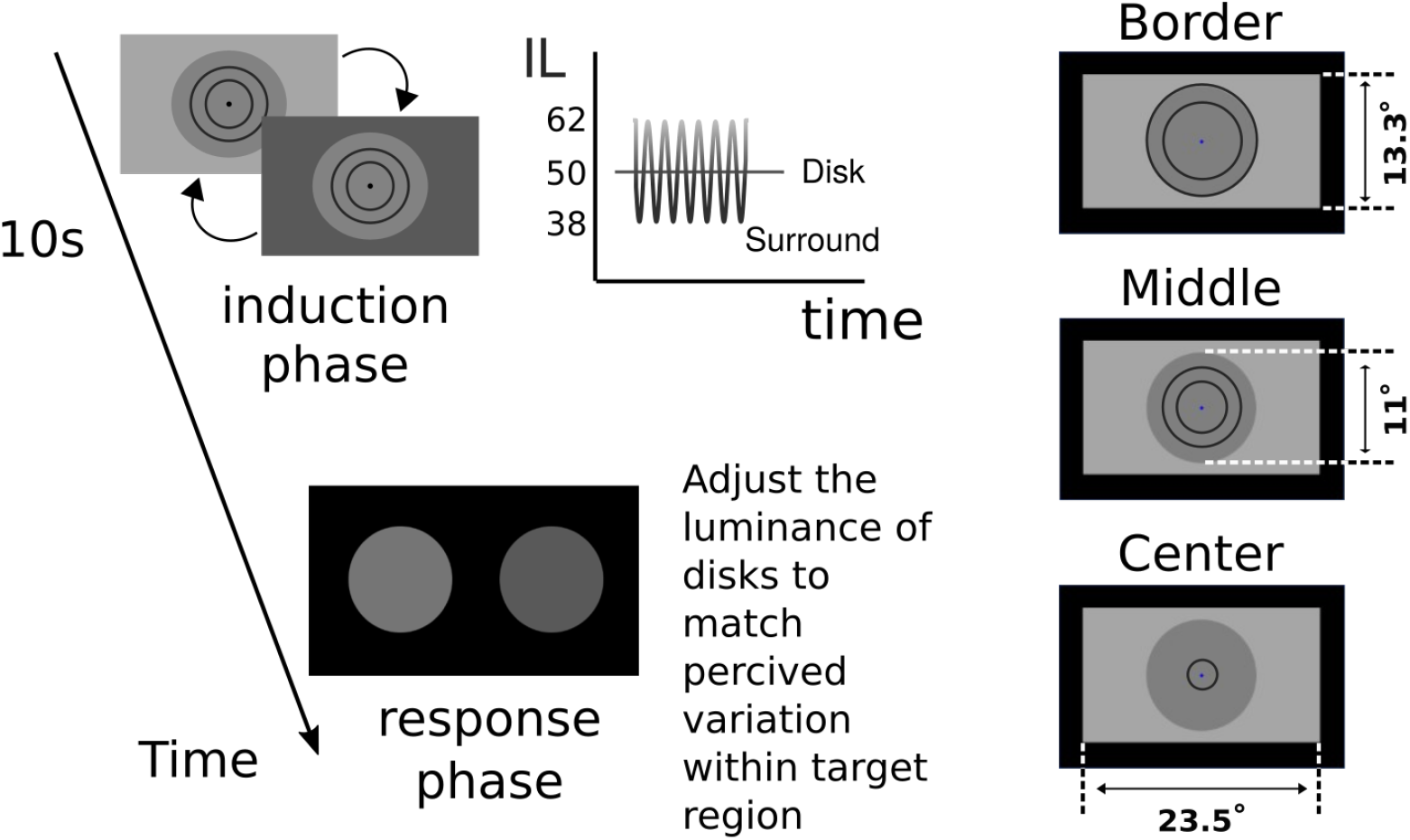
A trial in the behavioral experiment was comprised of an induction and a response phase. During the induction phase, we presented the dynamic SLI stimulus at different surround-modulation frequencies ranging from 0.25 to 16 Hz. Participants were instructed to fixate on the fixation mark and judge the perceived lightness variation in a target region marked by thin concentring circles. The right column shows the three target regions, border (8°-11°), middle (4°-7°), and center (0°-3°).

where *Lmin* and *Lmax* are the minimum and maximum luminance values of the screen, respectively, and *L* is the luminance of the stimulus (disk or surround). The luminance of the disk was kept constant at 50% IL, and the luminance of the surround varied between 62% and 38% IL, as described in more detail in the next section.

#### Procedure

An experimental session started with a three-second fixation period during which the luminances of the disk (50% IL) and surround (62% IL) were constant. After this, experimental trials started. Each trial consisted of an induction and a response phase. During the induction phase, the luminance of the disk remained constant (50% IL), and the surround luminance was sinusoidally modulated between 62% (starting value) and 38% IL, with one of 8 frequencies, 0.25, 0.5, 1, 2, 4, 6, 8, and 16 Hz. This produced a perceptual modulation in the disk out of phase with surround modulation, particularly for the lower frequencies. During both the fixation and the induction periods, participants were asked to fixate on a small blue mark at the center of the disk. In each trial, participants judged the perceived luminance variation within one of three target regions of the disk, center region (0°-3°), middle region (4°-7°), and border region (8°-11°).

After the induction phase ended, the response phase started, during which two luminance-adjustable disks were presented side by side on a black background. Participants were asked to adjust the luminance of the disks such that the difference between them corresponds to the difference between the perceived maximum and minimum luminance of the target region within the disk in the stimulation phase. The luminance of the disks could be adjusted by pressing the up and down arrow keys, and the trial was finalized by pressing the space bar, after which the next trial started. The luminance difference between the two disks was taken as the magnitude of SLI.

Overall, there were 24 conditions (8 frequency x 3 target regions), and each condition was repeated ten times, making a total of 240 trials. Trials were randomized across participants. Data were acquired in 3 sessions from each participant.

#### Behavioral data analysis

To analyze the behavioral data, we ran Linear Mixed Models (LMMs) using the ‘lme4’ package (Bates, Mächler, Bolker, & Walker, 2015) in RStudio (RStudio Team, 2020). The magnitude of the dynamic simultaneous lightness induction (SLI) in units of percent instrument luminance (% IL) was taken as the dependent variable. Eccentricity, frequency, and their interactions were included as fixed effects, and the between-participant variance was estimated using a random intercept in the model. The model was specified as

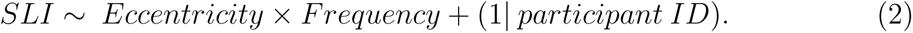

To calculate the significance of the fixed effects, we used the ‘lmerTest’ package (Kuznetsova, Brockhoff, & Christensen, 2017), which uses Satterthwaite’s method to estimate the degrees of freedom and generate p-values for linear mixed models. The assumptions of linear mixed models were met: The normality assumption of residuals was assessed by examination of the QQ-plot and Shapito-Wilk test, and the homogeneity of residual variance assumption was assessed by Levene’s test. The package ‘emmeans’ (Lenth, 2023) was used for obtaining the contrasts between estimated marginal means adjusted for multiple comparisons using Tukey’s test. The package ‘ggplot2’ was used for data visualization (Wickham, 2016).

### fMRI Experiment

#### Experimental Setup and Data Acquisition

A 3 Tesla Siemens Trio MR scanner (AG, Erlangen, Germany) with a 32-channel phase-array head coil was used to acquire the MR data at the National Magnetic Resonance Research Center (UMRAM), Bilkent University. Each participant underwent multiple MR runs in a session, including one run for high-resolution anatomical imaging (T1-weighted 3D MPRAGE sequence, spatial resolution: 1 mm^3^ isotropic, TR: 2600 ms, TE: 2.92 ms, flip angle: 12°, number of slices: 176), and a total of 8 functional runs, one for retinotopic mapping, one for functional region of interest (ROI) identification, and six for experimental conditions (T2*-weighted EPI sequence, spatial resolution: 3x3x3 mm^3^, TR: 2000 ms, TE: 35 ms, flip angle: 75°, number of slices: 30, slice orientation: parallel to calcarine sulcus). An MR-safe 32-inch LCD monitor (T-32, Troyka Med A.S., Ankara, Türkiye) was used to present the visual stimuli. The maximum and minimum achromatic luminance of the screen was 79.6 and 0.0024 cd/m^2^, respectively. The participants lay supine in the scanner and viewed the screen using a mirror attached to the head coil above their eyes, with a total viewing distance of about 172 cm.

#### MR Data Preprocessing and Analysis Software

MR data were pre-processed and analyzed using FSL and Freesurfer (Fischl, 2012; Jenkinson, Beckmann, Behrens, Woolrich, & Smith, 2012). Preprocessing steps included BET brain extraction applied on T1 anatomical images to remove non-brain tissue. For functional T2* images, preprocessing steps included MCFLIRT motion correction, BET brain extraction, and high-pass temporal filtering. Functional images were aligned to the individual’s T1 anatomical image and registered to standard T1 MNI152 brain resampled at 1 mm cubic voxels. For each individual participant, the 3D cortical surface was constructed from anatomical images using Freesurfer for visualizing statistical maps, delineating early visual areas, and identifying regions of interest.

#### Retinotopic Mapping and ROI identification

Horizontal and vertical wedges mapped with flickering checkerboard patterns were used in a block-design fMRI run to delineate the early visual areas V1, V2, V3, and V4 of each participant following the standard protocols in literature (Engel et al., 1997; Sereno et al., 1995). Stimulation code was written in the Java programming platform by us. The data were preprocessed as described above, and a general linear model (GLM) was applied using FSL. The resulting parametric maps were projected onto the 3D reconstruction of the participant’s cortex in native space. Then, Freesurfer’s drawing tools on its Freeview program were used to draw the boundaries between the early visual areas.

ROIs were identified within each visual area using a block-design localizer run. Stimulation code was written by us using the Psychtoolbox libraries (Brainard, 1997) on MATLAB (version 2018a, MathWorks Inc., New York, NY, USA). The basic stimuli were pairs of arcs mapped with flickering high-contrast checkerboard patterns (Figure 4). During the localizer run, in each 16-sec block, one of four pairs of arcs was presented to the participants at different eccentricities. The positions of the arcs corresponded to the center (1.5 to 2.5 degrees eccentricity), middle (5 to 6.5 degrees eccentricity), border (9 to 10.5 degrees eccentricity), and surround (14 to 16 degrees eccentricity) of the experimental disk. The data were preprocessed, and a general linear model (GLM) was applied using FSL. The resulting parametric maps were projected onto the 3D reconstruction of the participant’s cortex in native space. Based on this and the retinotopic mapping, ROIs within each early visual area were identified, e.g., V1 middle ROI, etc.

**Figure 4:**
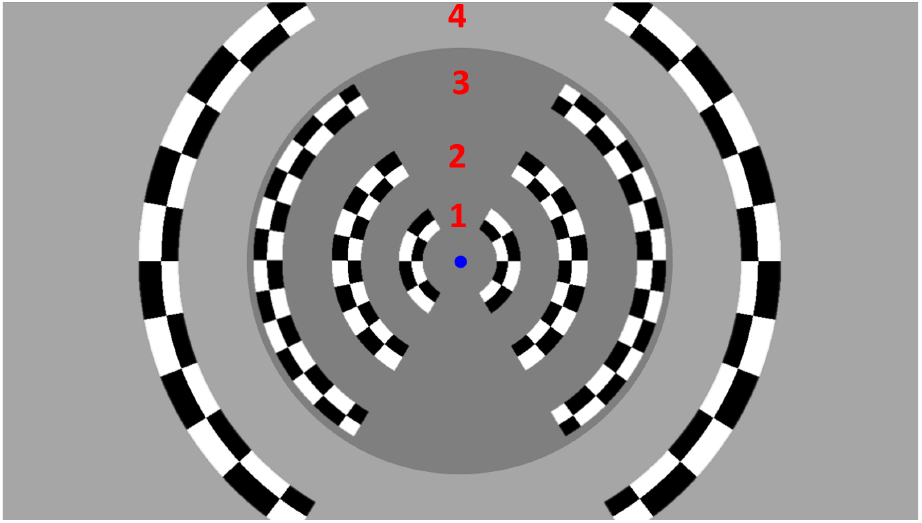
Functional localizer stimuli. High-contrast counter-phase flickering checker-board patterns were presented in pairs of arcs on a uniform gray background (arcs are overlaid on the experimental stimulus in this figure to provide a visual reference to the readers). Arcs 1, 2, 3, and 4 were used to localize the center (1.5°-2.5°), middle (5°-6.5°), border (9°-10.5°) and surround (14°-16°) ROIs in V1, V2, and V3. For V4, only border and center ROIs were extracted.

#### Experimental Design, Stimuli, and Data Analyses

fMRI responses to experimental stimuli were collected using a block-design paradigm. Three conditions, surround-modulation, disk-modulation, and control, were tested in different runs (Figure 2). There were two runs per condition. In all three conditions, the basic stimulus was composed of a gray-scale rectangle (surround) and a central disk with the same dimensions as in the behavioral experiment. In the surround-modulation condition, as in the behavioral experiment, the luminance of the disk was kept constant (27.6% IL), and the surround luminance was modulated sinusoidally (between 20.2% and 35.6% IL). In the disk-modulation condition, the luminance of the disk was sinusoidally modulated while the luminance of the surround was kept constant (27.6% IL). Here, the modulation of the disk followed the same temporal sinusoidal pattern as the surround in the surround-modulation condition (between 20.2% and 35.6% IL). The stimulus in the control condition was the same as in the surround-modulation condition, except that the disk was black (0% IL). Four sinusoidal modulation frequencies, 1, 2, 4, and 8 Hz, were tested in all conditions.

Each run started with a static block where a static screen composed of the disk, the surround, and the fixation mark was presented for a period of 12 seconds to allow for the fMRI signal to stabilize. This was followed by a 12-second test block in which the luminance of the surround (surround-modulation and control conditions) or the disk (disk-modulation condition) was modulated at one of the four modulation frequencies. The dynamic block was followed by another static block, and this pattern was repeated during the run such that each frequency condition was presented four times. Frequency conditions were pseudo-randomized and counter-balanced to rule out any order effects. Throughout the run, the fixation mark remained visible in all stimulation blocks. The default color of the fixation mark was blue, and it changed to red or yellow every 1500-2500 ms for 100 ms. Participants were asked to keep their fixation at the mark and report any changes in its color using an MR-safe response box (Fiber Optic Response Devices Package 904, Current Designs). The run was concluded with a 12-second static block. The total duration of an experimental run was 396 seconds.

Using FSL, we first ran a whole-brain GLM analysis. Using the functional ROIs as masks and the Featquerry tool of FSL, we extracted the GLM beta-weights from each ROI and treated them as fMRI response amplitudes. Further statistical analyses were performed on the extracted fMRI response amplitudes in RStudio (RStudio Team, 2020).

For V1, we ran Linear Mixed Models (LMMs) using the ‘lme4’ package (Bates et al., 2015) where the magnitude of the fMRI response amplitude was taken as the dependent variable. Condition (three levels; disk modulation, surround modulation, and control) Eccentricity (four levels; center, middle, border and surround), frequency (four levels; 1, 2, 4, and 8 Hz), and all the subsequent interactions were included as fixed effects. The between-participant variance was estimated using a random intercept in the model. The model was specified as

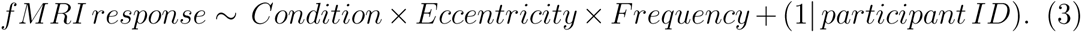

For each of V2, V3, and V4, a separate model was specified. We report only the results in the center ROIs because responses in those ROIs are least affected by the border contrast variations, as explained further in the Results section. Thus, eccentricity is not included as a fixed effect factor in the models. The rest of the specifications were the same as the model for V1:

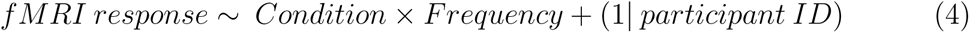

As in the behavioral analysis, we used the ‘lmerTest’ package (Kuznetsova et al., 2017) to calculate the significance of the fixed effects. We found that the assumptions of linear mixed models were met. The normality assumption of residuals was assessed by examination of QQ-plots and Shapito-Wilk test. The homogeneity of residual variance assumption was assessed by Levene’s test. The package ‘emmeans’ (Lenth, 2023) was used for obtaining the contrasts between estimated marginal means adjusted for multiple comparisons using Tukey’s test.

## Results

### Behavioral experiment

Figure 5 shows the means of the magnitude of the dynamic simultaneous lightness induction (SLI) as a function of surround modulation frequency. Consistent with previous literature, we found that SLI decreases with surround modulation frequency. This trend is observed in all eccentricities tested. Linear mixed model (LMM) analysis on the SLI magnitudes showed a significant main effect of frequency (F(7) = 116.3; p*<*0.001), a significant main effect of eccentricity (F(2) = 183.4713; p*<*0.001), as well as a significant interaction between eccentricity and frequency (F(14) = 3.0559; p*<*0.001). As can be observed in Figure 5, the induction is significantly stronger near the border between the disk and the surround. This is also reflected in post hoc pairwise comparisons of estimated marginal means among eccentricity conditions, which show that the magnitude of SLI is significantly different between all eccentricity conditions (p*<*0.01), with the border region producing the highest estimated marginal mean (18.56) in comparison to the middle (10.81) and center (9.22). These results show that for our stimulus configuration, lightness induction at the center of the disk is stronger for lower modulation frequencies, decreases for higher frequencies, and almost disappears at 8 and 16 Hz. Based on these results, we chose four surround modulation frequencies, 1, 2, 4, and 8 Hz, to be tested in the fMRI experiment.

**Figure 5:**
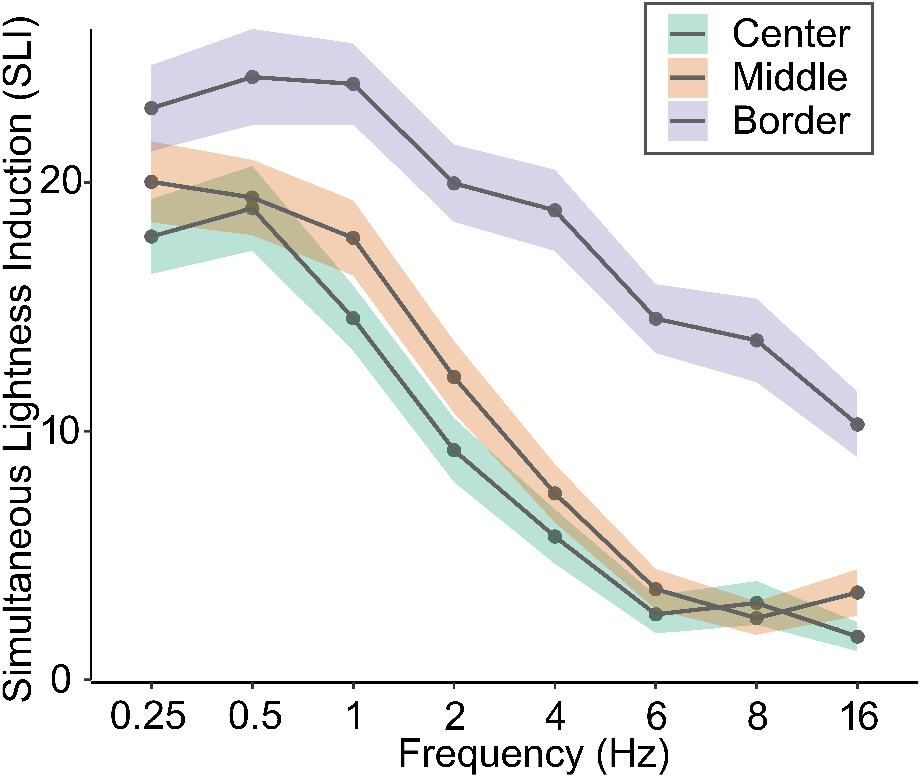
Behavioral results. Group average of the dynamic simultaneous lightness induction (SLI) as a function of surround modulation frequency for each eccentricity condition. SLI is higher at the outer eccentricity, which is closest to the border between the disk and the surround. In the middle and center eccentricities, SLI is highest at lower surround modulation frequencies, decreases with frequency, and nearly disappears for the highest frequencies. Shaded regions show the standard error around the mean.

### fMRI experiment

#### V1 Results

Figure 6 shows the fMRI responses from the primary visual cortex (V1) averaged over ten participants under the disk-modulation and surround-modulation conditions. Linear Mixed Models (LMMs) analyses with conditions (disk-modulation, surround-modulation, and control), frequency (1, 2, 4, and 8 Hz), and eccentricity (center, middle, border, and surround) as fixed effect factors revealed a significant main effect of eccentricity (F(3) = 28.3; p *<*0.001), a significant main effect of frequency (F(3) = 27.7 ; p *<*0.001), a significant main effect of condition (F(2) = 55.8 ; p *<*0.001), and a significant interaction between eccentricity and condition (F(6) = 8.2; p *<*0.001). The responses in the border ROI under the surround-modulation and disk-modulation conditions are not different (p=0.8352), and they both increase with frequency (Figure 6, Border ROI panel). This shows that the responses in the border ROI are dominated by the contrast variation at the border between the disk and the surround. We further note that the fMRI responses are smaller in the center and middle ROIs compared to the border ROI at all frequencies. This reduction appears systematic and depends on the distance between the border and the ROI. In light of these observations, we focus only on the center ROI in our analyses, as it is likely to be least affected by the dynamic border contrast and best represents the neural activity related to lightness processing.

**Figure 6:**
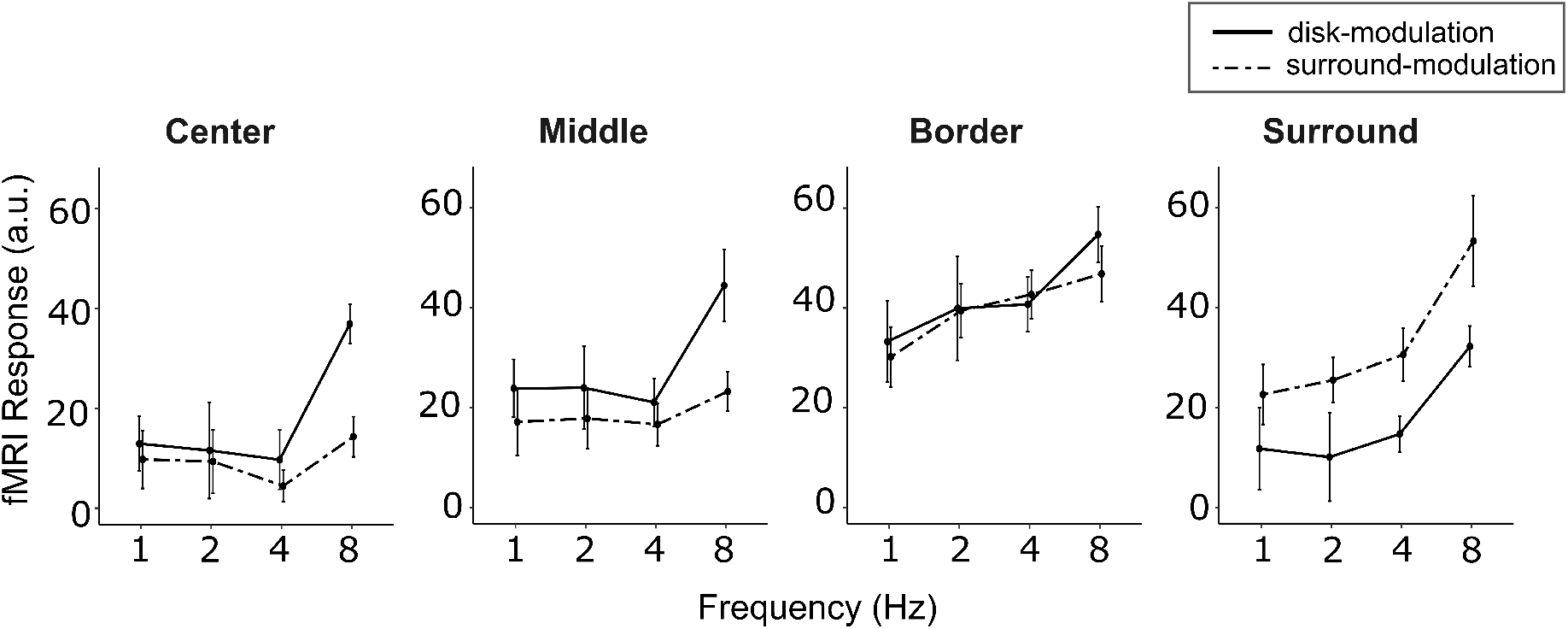
V1 group average of fMRI responses as a function of frequency in all four ROIs. fMRI responses near the border between the disk and surround are the same for the surround-modulation and disk-modulation conditions for all frequencies (Border ROI plot). This shows that those responses are likely caused by the contrast modulation at the border. Whereas in the central and middle ROIs, fMRI responses under the surround-modulation and disk-modulation conditions remain similar only at 1, 2, and 4Hz, but diverge at 8Hz (Central ROI and Middle ROI plots), supporting that V1 responses in those ROIs are correlated with SLI, not simply the contrast modulation at the border. Error bars: SEM.

Figure 7 left panel shows averaged fMRI responses in the center ROI. Posthoc pairwise comparison of estimated marginal means from LMM revealed a significant difference between the surround-modulation and disk-modulation conditions only at 8 Hz (p *<*0.05). To further aid visualization, we computed the differences between the responses under the two conditions (Figure 7, right panel). Those results show that relative responses to surround modulation, just as the behavioral SLI effect, decrease with frequency. This provides evidence that the responses reflect neural activity related to the context-dependent SLI effect.

**Figure 7:**
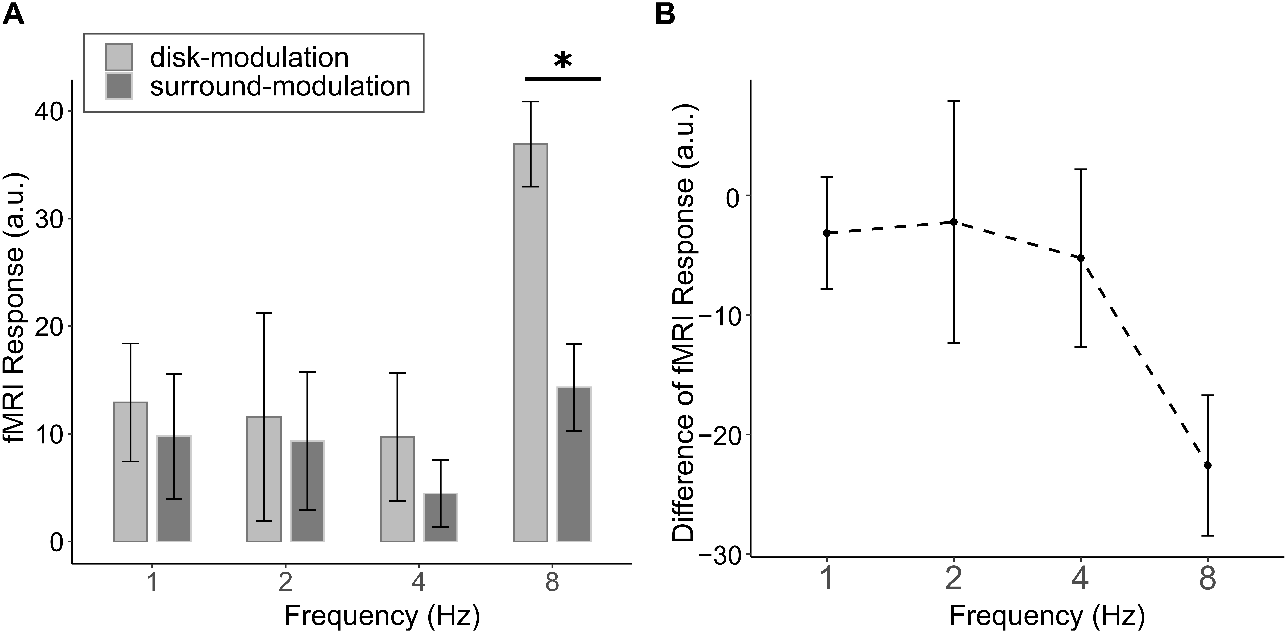
**A**. fMRI responses in the center ROI under surround-modulation and disk-modulation conditions, replotted from Figure 6 center ROI plot. * indicate a significant difference under the pairwise comparison of estimated marginal means. **B**. The difference between responses under the two conditions, surround-modulation minus disk-modulation, as a function of frequency.

Figure 8 shows the fMRI responses under the control condition in the center ROI, together with the responses under the surround-modulation and disk-modulation conditions. At 1 and 2 Hz, fMRI responses under the surround-modulation condition are stronger than those under the control condition and similar to those under the disk-modulation condition. But at higher frequencies, particularly at 8 Hz, fMRI responses under the surround-modulation condition are close to those under the control condition and less than those under the disk-modulation condition. These results further support that fMRI responses under the surround-modulation condition reflect neural activity related to perceptual lightness induction, not long-range luminance responses (as described in Figure 2).

**Figure 8:**
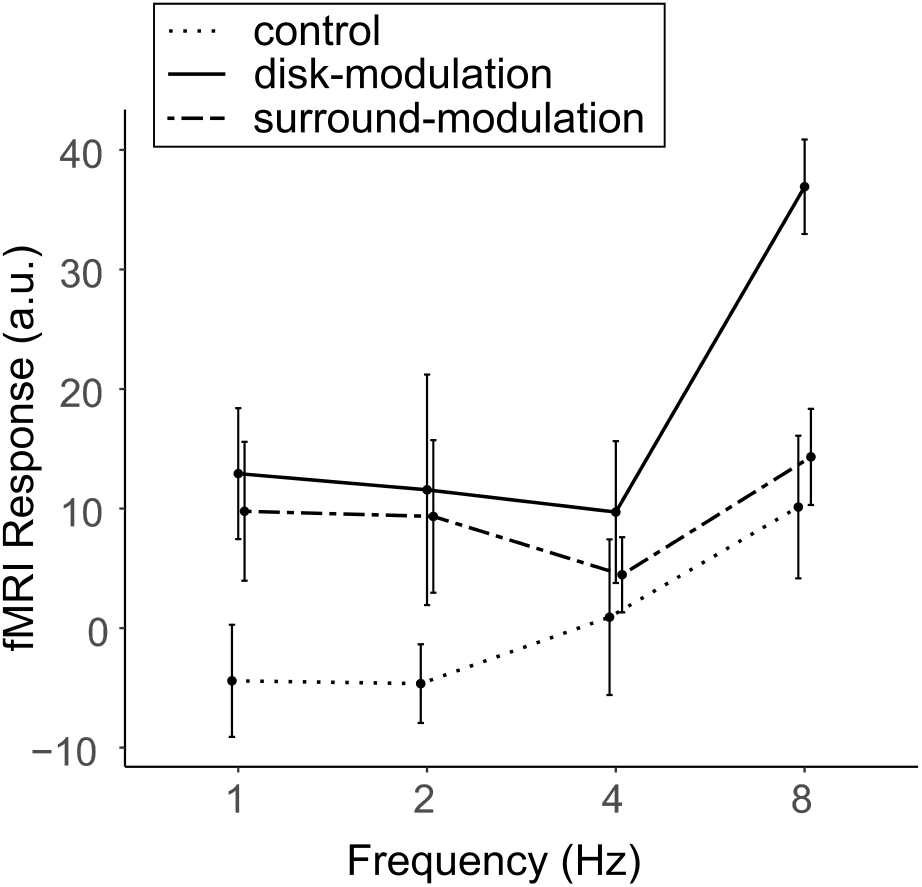
fMRI responses in the V1 center ROI under the surround-modulation, disk-modulation, and control conditions. At low frequencies, responses under the surround-modulation condition are approximately the same as those under the disk-modulation condition and larger than those under the control condition. Whereas, at high frequencies, especially at 8Hz, this relation reverses. At 8 Hz, the responses under the surround-modulation condition are approximately the same as those under the control condition and lower than those under the disk-modulation condition. This is exactly what would be expected from an area whose activity is correlated with the behavioral lightness induction effect, as depicted in Figure 2. Error bars represent SEM.

#### Extrastriate Areas

Figure 9 shows averaged fMRI responses in the center ROIs of visual areas V2, V3, and V4. Linear Mixed Models (LMMs) analysis, which was performed on each visual area separately, with conditions (disk-modulation, surround-modulation, and control) and frequency (1, 2, 4, and 8 Hz) as fixed effect factors, revealed a significant main effect of condition in all areas (p *<*0.001), a main effect of frequency only in V4 (p *<*0.05), and no significant interactions between condition and frequency. Posthoc pairwise comparison of estimated marginal means revealed that fMRI responses under the control condition are significantly different from those under the disk-modulation and surround-modulation conditions. However, there was no significant difference between surround-modulation and disk-modulation conditions in any area. This pattern of results resembles the last possible outcome depicted in Figure 2, and suggests that the responses are dominated by contrast variation at the border, thus, it does not provide strong evidence that V2, V3, and V4 are involved in context-dependent lightness processing. We discuss these findings in the next section.

**Figure 9:**
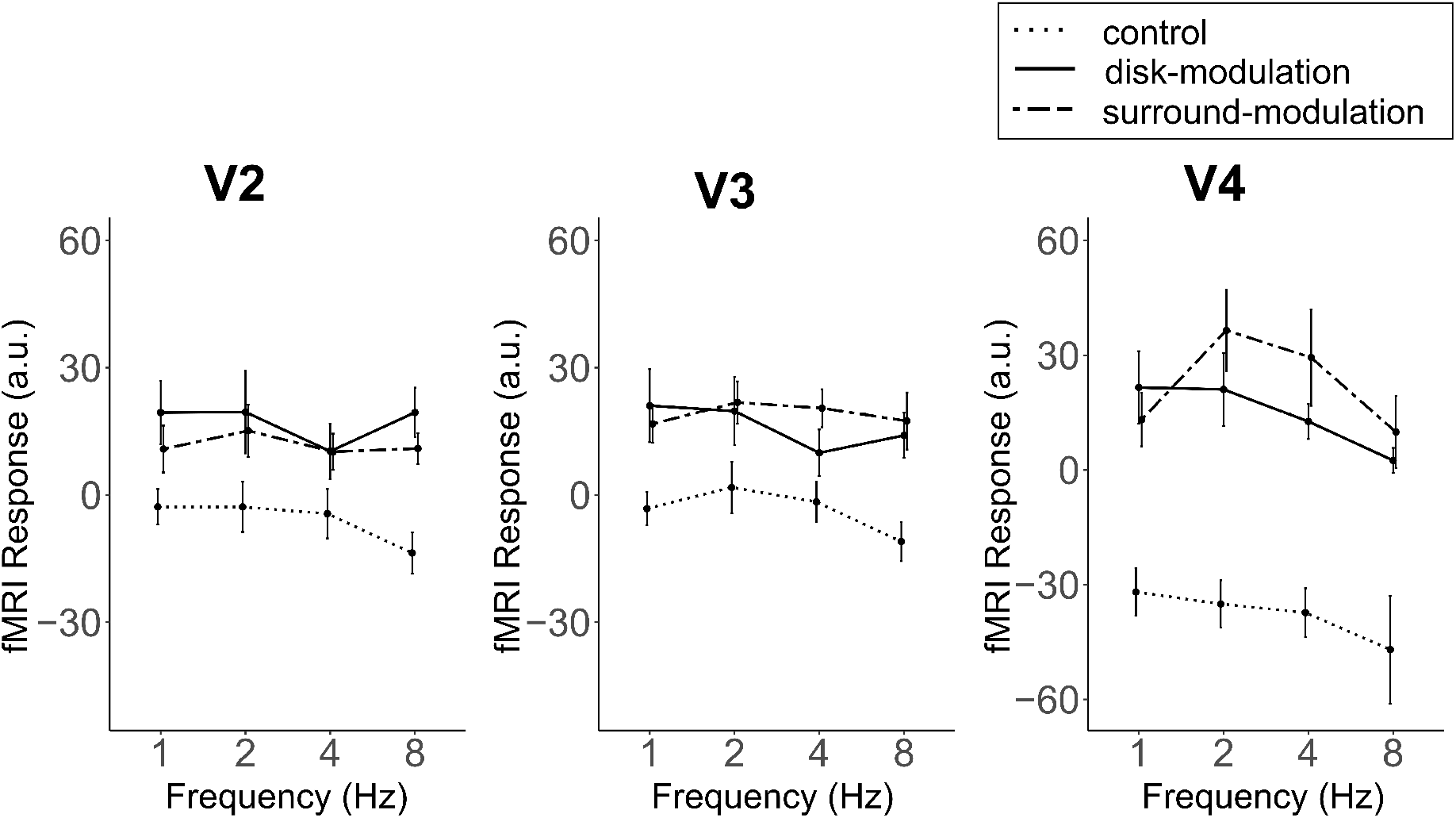
fMRI responses in V2, V3, and V4 center ROIs. Because it was not possible to localize the center ROIs in all participants, the data were averaged across 8 participants in V2 and V3 and across 7 participants in V4. Responses under the disk-modulation and surround-modulation conditions did not differ in any area, and they were larger than those under the control condition. The pattern of results suggests that the fMRI response is dominated by contrast variations at the border, as depicted in Figure 2. Error bars represent SEM.

## Discussion

Our results show that fMRI responses in the primary visual cortex (V1) correlate with perceived lightness in the dynamic simultaneous lightness induction (SLI) effect. This outcome supports that V1 is involved in context-dependent lightness processing. We did not, however, find strong fMRI evidence that extrastriate visual areas, V2, V3, and V4, are involved in context-dependent lightness processing.

First, in a behavioral experiment, we tested the strength of SLI. Consistent with the findings in the literature, we found that the dynamic SLI effect is stronger at lower surround modulation frequencies, decreases with increasing frequency, and nearly vanishes at very high frequencies (De Valois et al., 1986; Pereverzeva & Bromfield, 2013; Rossi & Paradiso, 1996). Next, in an fMRI experiment, we measured responses from visual areas while observers viewed the dynamic SLI stimulus. We found that V1 responses in the region corresponding to the center of the disk reflect the pattern in the behavioral results: lightness-related responses are stronger at low frequencies than those at high frequencies. This finding is in line with previous animal electrophysiological studies (Kinoshita & Komatsu, 2001; MacEvoy & Paradiso, 2001; Rossi, Desimone, & Ungerleider, 2001; Rossi & Paradiso, 1999; Rossi et al., 1996) as well as human fMRI studies (Boyaci et al., 2007, 2010; Haynes et al., 2004; Pereverzeva & Murray, 2008; Sasaki & Watanabe, 2004), and support that the activity of V1 correlates with context-dependent lightness perception.

To address the possible confounding effect of border contrast variations (e.g., see Cornelissen et al. 2006), we systematically assessed how the fMRI responses varied as the modulation frequency increased. We performed this analysis in functionally defined regions of interest (ROIs) corresponding to different parts of the disk. If the responses were primarily driven by the border contrast, not induced lightness, then we would expect them to increase with frequency (Chai et al., 2019; Goodyear & Menon, 1998). But that was not the case for all ROIs. In the ROI closest to the border of the disk (border ROI), the responses indeed increased with frequency, indicating that they are largely driven by the border contrast. fMRI responses to the parts of the disk further from the border (central and middle ROIs), however, showed little or no increase with frequency. This pattern speaks against a border contrast explanation.

To further assess how activity varies with frequency, we computed the responses under the surround-modulation condition relative to those under the disk-modulation condition, where the surround remains static, and the disk is modulated in luminance. Because the border contrast variation was the same across the two conditions, this analysis allowed us to compare the responses to luminance and induced lightness variations directly. At low frequencies, where there is a strong behavioral effect, the responses under the surround-modulation and disk-modulation conditions were nearly equal in the ROI corresponding to the center of the disk. But at the highest frequency, where induction disappears, the responses to the surround-modulation condition were significantly less than those to the disk-modulation condition. In other words, relative responses under the surround-modulation condition decreased with frequency, in line with the behavioral results (Figure 7).

We noted that the fMRI responses under the surround-modulation condition might not be related to context-dependent lightness but merely reflect the activity of neurons that respond to the surround modulation or the overall illumination in the environment. A comparison of responses under the surround-modulation and control conditions rules out this possibility. Under both conditions, the surround is modulated with the same temporal characteristics. But only under the surround-modulation condition does the disk’s lightness vary. If the fMRI responses merely reflect the neural activity in response to the surround modulation, they should be the same under both conditions at all frequencies. But this is not the case. At the highest frequency, where there is no lightness variation under the surround-modulation condition, fMRI responses are almost equal under the surround-modulation and control conditions (Figure 8). This suggests that when perceptually there is no induction, the fMRI response might indeed be driven by the surround variation alone. But at low frequencies, the fMRI responses under the surround-modulation condition are stronger than those under the control condition. In other words, when there is lightness induction, the fMRI responses are stronger compared to when there is no lightness induction. This pattern argues against a long-range response explanation.

### Extrastriate Areas

Previous studies have found evidence that extrastriate areas V2, V3, and V4 are involved in context-dependent lightness processing (Boyaci et al., 2007, 2010; Bushnell, Harding, Kosai, Bair, & Pasupathy, 2011; Hung, Ramsden, & Roe, 2007; Perna et al., 2005; Roe, Lu, & Hung, 2005; Ruff, Brainard, & Cohen, 2018; Salmela & Vanni, 2013; van de Ven, Jans, Goebel, & De Weerd, 2012; Zhou et al., 2020). Here we found that, in V2, V3, and V4 center ROIs, fMRI responses under the disk-modulation and surround modulation conditions were nearly the same, and they were both larger than those under the control condition at both low and high frequencies. At low frequencies, this is the expected pattern of results in an area involved in lightness processing. The pattern at high frequencies, however, suggests that the responses may be caused by the border contrast variation and raises doubts that they are related to the lightness induction (the last possible outcome depicted in Figure 2). Thus, our results provide limited support that extrastriate areas V2, V3, and V4 are involved in context-dependent lightness processing.

This outcome could be the result of fMRI’s limitation as a technique to resolve neural activity. Specifically, a measurement from an fMRI unit, voxel, reflects an aggregate and indirect response from hundreds of thousands of neurons that may have heteroge-neous response characteristics. One of those characteristics is the (population) receptive field size, which is typically larger in extrastriate areas compared to V1 (Smith, Singh, Williams, & Greenlee, 2001; van Dijk, de Haas, Moutsiana, & Schwarzkopf, 2016). Even though we have carefully identified voxels that, on average, respond more strongly to our flickering checkerboard localizer near the center of the disk, these voxels may still harbor neurons that have relatively large RF sizes and respond to the contrast variation at the border. Subtle variations with frequency in the lightness-related neural activity could be buried under those long-range activations. Thus, our fMRI results do not unequivocally prove that extrastriate areas are not involved in context-dependent lightness processing.

Interestingly, unlike in V1, fMRI responses in extrastriate areas do not increase with increasing frequency under the disk-modulation condition. In V4, we even observe a decrease in responses with frequency. Previous fMRI studies investigating responses of the visual cortex to chromatic or/and achromatic temporal modulations reported that V4 shows a low pass temporal dependence on frequency (D’Souza, Auer, Strasburger, Frahm, & Lee, 2011; Mullen, Thompson, & Hess, 2010). This reduced temporal resolution in V4 can be attributed to its involvement in form perception and object recognition, where temporal information may play a less critical role. Further, V4 plays a role in surface perception, and its responses may show dissociation with those of V1 if the stimulus does not appear to be a surface (Bouvier et al., 2008). Thus, an alternative explanation for our results could be that at high frequencies, V4 may not treat the stimulus as a surface anymore, leading to a decrease in its responses with frequency.

## Conclusion

Our findings show that the fMRI responses in V1 correlate with dynamic lightness induction and thus provide further evidence that V1 is involved in context-dependent lightness processing. We did not find, however, strong fMRI evidence that extrastriate areas, V2, V3, and V4, are also involved in context-dependent lightness processing.

## References

Adelson, E. H. (1993). Perceptual organization and the judgment of brightness. Science, 262 (5142), 2042–2044.

Alhazen, I. (1883). Book of optics (a. sabra, trans.). The otics of Ibn al-Haytham, 2.

Anderson, E. J., Dakin, S. C., & Rees, G. (2009). Monocular signals in human lateral geniculate nucleus reflect the craik–cornsweet–o’brien effect. Journal of Vision, 9 (12), 14–14.

Bates, D., Mächler, M., Bolker, B., & Walker, S. (2015). Fitting linear mixed-effects models using lme4. Journal of Statistical Software, 67 (1), 1–48. doi: 10.18637/jss.v067.i01

Blakeslee, B., & McCourt, M. E. (1997). Similar mechanisms underlie simultaneous brightness contrast and grating induction. Vision research, 37 (20), 2849–2869.

Blakeslee, B., & McCourt, M. E. (1999). A multiscale spatial filtering account of the white effect, simultaneous brightness contrast and grating induction. Vision research, 39 (26), 4361–4377.

Bouvier, S. E., Cardinal, K. S., & Engel, S. A. (2008). Activity in visual area v4 correlates with surface perception. Journal of vision, 8 (7), 28–28.

Boyaci, H., Fang, F., Murray, S. O., & Kersten, D. (2007). Responses to lightness variations in early human visual cortex. Current Biology, 17 (11), 989–993.

Boyaci, H., Fang, F., Murray, S. O., & Kersten, D. (2010). Perceptual grouping-dependent lightness processing in human early visual cortex. Journal of vision, 10 (9), 4–4.

Brainard, D. H. (1997). The psychophysics toolbox. Spatial vision, 10 (4), 433–436.

Bushnell, B. N., Harding, P. J., Kosai, Y., Bair, W., & Pasupathy, A. (2011). Equiluminance cells in visual cortical area v4. Journal of Neuroscience, 31 (35), 12398–12412.

Chai, Y., Handwerker, D. A., Marrett, S., Gonzalez-Castillo, J., Merriam, E. P., Hall, A., . . . Bandettini, P. A. (2019). Visual temporal frequency preference shows a distinct cortical architecture using fmri. Neuroimage, 197, 13–23.

Cornelissen, F. W., Wade, A. R., Vladusich, T., Dougherty, R. F., & Wandell, B. A. (2006). No functional magnetic resonance imaging evidence for brightness and color filling-in in early human visual cortex. Journal of Neuroscience, 26 (14), 3634–3641.

Dakin, S. C., & Bex, P. J. (2003). Natural image statistics mediate brightness ‘filling in’. Proceedings of the Royal Society of London. Series B: Biological Sciences, 270 (1531), 2341–2348.

De Valois, R. L., Webster, M. A., De Valois, K. K., & Lingelbach, B. (1986). Temporal properties of brightness and color induction. Vision research, 26 (6), 887–897.

D’Souza, D. V., Auer, T., Strasburger, H., Frahm, J., & Lee, B. B. (2011). Temporal frequency and chromatic processing in humans: an fmri study of the cortical visual areas. Journal of Vision, 11 (8), 8–8.

Economou, E., Zdravkovic, S., & Gilchrist, A. (2007). Anchoring versus spatial filtering accounts of simultaneous lightness contrast. Journal of vision, 7 (12), 2–2.

Fischl, B. (2012). Freesurfer. Neuroimage, 62 (2), 774–781.

Geisler, W. S., Albrecht, D. G., & Crane, A. M. (2007). Responses of neurons in primary visual cortex to transient changes in local contrast and luminance. Journal of Neuroscience, 27 (19), 5063–5067.

Gilchrist, A., Kossyfidis, C., Bonato, F., Agostini, T., Cataliotti, J., Li, X., . . . Economou, E. (1999). An anchoring theory of lightness perception. Psychological review, 106 (4), 795.

Goodyear, B. G., & Menon, R. S. (1998). Effect of luminance contrast on bold fmri response in human primary visual areas. Journal of Neurophysiology, 79 (4), 2204–2207.

Haynes, J.-D., Lotto, R. B., & Rees, G. (2004). Responses of human visual cortex to uniform surfaces. Proceedings of the National Academy of Sciences, 101 (12), 4286–4291.

Heinemann, E. G. (1955). Simultaneous brightness induction as a function of inducing- and test-field luminances. Journal of experimental psychology, 50 (2), 89.

Hung, C. P., Ramsden, B. M., & Roe, A. W. (2007). A functional circuitry for edge-induced brightness perception. Nature neuroscience, 10 (9), 1185–1190.

Jenkinson, M., Beckmann, C. F., Behrens, T. E., Woolrich, M. W., & Smith, S. M. (2012). Fsl. Neuroimage, 62 (2), 782–790.

Kingdom, F. (1997). Simultaneous contrast: the legacies of hering and helmholtz (Vol. 26) (No. 6). SAGE Publications Sage UK: London, England.

Kinoshita, M., & Komatsu, H. (2001). Neural representation of the luminance and brightness of a uniform surface in the macaque primary visual cortex. Journal of neurophysiology, 86 (5), 2559–2570.

Komatsu, H. (2006). The neural mechanisms of perceptual filling-in. Nature reviews neuroscience, 7 (3), 220–231.

Kuznetsova, A., Brockhoff, P. B., & Christensen, R. H. B. (2017). lmerTest package: Tests in linear mixed effects models. Journal of Statistical Software, 82 (13), 1–26. doi: 10.18637/jss.v082.i13

Lenth, R. V. (2023). emmeans: Estimated marginal means, aka least-squares means [Computer software manual]. Retrieved from https://CRAN.R-project.org/package=emmeans (R package version 1.8.4–1)

MacEvoy, S. P., & Paradiso, M. A. (2001). Lightness constancy in primary visual cortex. Proceedings of the National Academy of Sciences, 98 (15), 8827–8831.

Mullen, K. T., Thompson, B., & Hess, R. F. (2010). Responses of the human visual cortex and lgn to achromatic and chromatic temporal modulations: an fmri study. Journal of vision, 10 (13), 13–13.

Pereverzeva, M., & Bromfield, W. D. (2013). Spatial nonuniformities and velocity of filling-in in dynamic brightness induction. Journal of Vision, 13 (5), 17–17.

Pereverzeva, M., & Murray, S. O. (2008). Neural activity in human v1 correlates with dynamic lightness induction. Journal of Vision, 8 (15), 8–8.

Perna, A., Tosetti, M., Montanaro, D., & Morrone, M. C. (2005). Neuronal mechanisms for illusory brightness perception in humans. Neuron, 47 (5), 645–651.

Roe, A. W., Lu, H. D., & Hung, C. P. (2005). Cortical processing of a brightness illusion. Proceedings of the National Academy of Sciences, 102 (10), 3869–3874.

Rossi, A. F., Desimone, R., & Ungerleider, L. G. (2001). Contextual modulation in primary visual cortex of macaques. Journal of Neuroscience, 21 (5), 1698–1709.

Rossi, A. F., & Paradiso, M. A. (1996). Temporal limits of brightness induction and mechanisms of brightness perception. Vision research, 36 (10), 1391–1398.

Rossi, A. F., & Paradiso, M. A. (1999). Neural correlates of perceived brightness in the retina, lateral geniculate nucleus, and striate cortex. Journal of Neuroscience, 19 (14), 6145–6156.

Rossi, A. F., Rittenhouse, C. D., & Paradiso, M. A. (1996). The representation of brightness in primary visual cortex. Science, 273 (5278), 1104–1107.

RStudio Team. (2020). Rstudio: Integrated development environment for r [Computer software manual]. Boston, MA. Retrieved from http://www.rstudio.com/

Ruff, D. A., Brainard, D. H., & Cohen, M. R. (2018). Neuronal population mechanisms of lightness perception. Journal of Neurophysiology, 120 (5), 2296–2310.

Salmela, V. R., & Vanni, S. (2013). Brightness and transparency in the early visual cortex. Journal of vision, 13 (7), 16–16.

Sasaki, Y., & Watanabe, T. (2004). The primary visual cortex fills in color. Proceedings of the National Academy of Sciences, 101 (52), 18251–18256.

Sinha, P., Crucilla, S., Gandhi, T., Rose, D., Singh, A., Ganesh, S., . . . Bex, P. (2020). Mechanisms underlying simultaneous brightness contrast: early and innate. Vision research, 173, 41–49.

Smith, A. T., Singh, K. D., Williams, A. L., & Greenlee, M. W. (2001). Estimating receptive field size from fmri data in human striate and extrastriate visual cortex. Cerebral cortex, 11 (12), 1182–1190.

van de Ven, V., Jans, B., Goebel, R., & De Weerd, P. (2012). Early human visual cortex encodes surface brightness induced by dynamic context. Journal of Cognitive Neuroscience, 24 (2), 367–377.

van Dijk, J. A., de Haas, B., Moutsiana, C., & Schwarzkopf, D. S. (2016). Intersession reliability of population receptive field estimates. NeuroImage, 143, 293–303. doi: 10.1016/j.neuroimage.2016.09.013

Vinke, L. N., & Ling, S. (2020). Luminance potentiates human visuocortical responses. Journal of Neurophysiology, 123 (2), 473–483.

Von Helmholtz, H. (1867). Handbuch der physiologischen optik (Vol. 9). Voss.

Wickham, H. (2016). ggplot2: Elegant graphics for data analysis. Springer-Verlag New York. Retrieved from https://ggplot2.tidyverse.org

Zhou, H., Davidson, M., Kok, P., McCurdy, L. Y., de Lange, F. P., Lau, H., & Sandberg, K. (2020). Spatiotemporal dynamics of brightness coding in human visual cortex revealed by the temporal context effect. Neuroimage, 205, 116277.

